# Association of inflammation and tissue damage induced biological processes in masseter muscle with the resolution of chronic myalgia

**DOI:** 10.1101/2023.04.21.537828

**Authors:** Karen A. Lindquist, Sergey A. Shein, Anahit H. Hovhannisyan, Jennifer Mecklenburg, Yi Zou, Zhao Lai, Alexei V. Tumanov, Armen N. Akopian

## Abstract

Biological processes linked to intramuscular inflammation during myogenous temporomandibular disorder (TMDM) are largely unknown. We mimicked this inflammation by intra-masseteric muscle (MM) injections of complete Freund’s adjuvant (CFA) or collagenase type 2 (Col), which emulates tissue damage. CFA triggered mechanical hypersensitivity at 1d post-injection was mainly linked to processes controlling chemotactic activity of monocytes and neutrophils. At 5d post-CFA, when hypersensitivity was resolved, there was minimal inflammation whereas tissue repair processes were pronounced. Low dose Col (0.2U) also produced acute orofacial hypersensitivity that was linked to tissue repair, but not inflammatory processes. High dose Col (10U) triggered prolonged orofacial hypersensitivity with inflammatory processes dominating at 1d post-injection. At pre-resolution time point (6d), tissue repair processes were underway and a significant increase in pro-inflammatory gene expressions compared to 1d post-injection were detected. RNA-seq and flow cytometry showed that immune processes in MM were linked to accumulation of macrophages, natural killer and natural killer T cells, dendritic cells and T-cells. Altogether, CFA and Col treatments induced different immune processes in MM. Importantly, orofacial hypersensitivity resolution was preceded with repairs of muscle cell and extracellular matrix, an elevation in immune system gene expression and accumulation of distinct immune cells in MM.

## Introduction

Myogenous temporomandibular disorder (TMDM)^1^ is one of most prevalent types of myofascial pain syndromes^2–4^. Chronic TMDM affects approximately 10% of the population, with two-thirds being women^2^. Precise etiology, pathogenesis, and pathophysiology underlying chronic TMDM remains unclear. Nevertheless, studies showed that TMDM is often accompanied by myositis with mild inflammation^1, 5^. Individual inflammatory mediators, such as TNF-a, IL-1β, IL-6 and IL-8, in the masseter muscle (MM) were revealed in TMDM patients using a variety of approaches including micro-dialysis^1, 5–8^. However, the overall biological processes in MM linked to this mild inflammation is largely unknown.

TMDM is thought to develop due to muscle ischemia following consistent and repetitive physical overloading of masticatory muscles^1, 5^. Muscle overload can disrupt local blood flow to the muscles of mastication, triggering ischemia, myositis and inflammation ^9, 10^. Intramuscular inflammation can be mimicked by the standard approach of injecting complete Freund’s adjuvant (CFA) into the MM ^11, 12^. Inflammation caused by tissue damage differs from that produced by CFA^13^, and ischemia-induced muscle damage may involve extracellular matrix dysfunction/damage^14–16^ and collagen degradation^17, 18^. Therefore, we developed a model of intramuscular tissue damage by disrupting extracellular matrix using the matrix metalloproteinase collagenase type 2 (Col).

This study aimed to bridge a gap in knowledge related to the biological processes in different inflammatory conditions in MM induced by injections of CFA, low dose (0.2U) Col, or high dose (10U) Col mimicking tissue damage. MM tissues were collected after mechanical threshold measurements in mice at different time points. We then used bulk RNA seq to trace biological processes within inflamed MM. Additionally, certain data obtained from bulk RNA-seq were validated using flow cytometry and immunohistochemistry, which characterize the immune cell profiles in inflamed MM tissue.

## Materials and Methods

### Ethical Approval

All animal experiments conformed to protocols approved by the University Texas Health Science Center at San Antonio (UTHSCSA) Institutional Animal Care and Use Committee (IACUC). We followed guidelines issued by the National Institutes of Health (NIH) and the Society for Neuroscience (SfN) to minimize the number of animals used and their suffering. Mice were housed under controlled conditions (≈22°C), relative humidity 40-60%, and a 12-h light-dark cycle with lights on at 7:00 AM. Food and water were available *ad libitum* in their home cages.

### Key reagents and mouse lines

Experiments were conducted on wild-type adult (2-4-months-old) C57Bl/6 male mice, which were purchased from Jackson Laboratory (Bar Harbor, ME). The Ccr2^RFP^/Cx3cr1^GFP^ (Stock No: 032127) mouse line on the B6.129 background was purchased from the Jackson Laboratory (Bar Harbor, ME). Crude collagenase (Col) type 2 (>125 units per mg; Cat: LS004214) preparations were acquired from Worthington (Lakewood, NJ) and complete Freund’s adjuvant (CFA; Cat: F5881) was obtained from Millipore-Sigma (St. Louis, MO).

### Masseter muscle (MM) injection

MM injections with 10μl collagenase type 2 or CFA (1:1 = CFA : PBS) were performed under isoflurane anesthesia. The area of injection was swabbed with 70% alcohol beforehand. The site for injection was identified by palpating the zygomatic arch and the body of the mandible. A 30-gauge needle was inserted into the point inferior to the posterior third of the zygomatic arch and midway between the zygomatic arch and the body of the mandible. The needle was advanced in an anterior direction in about 3 mm. After a gentle aspiration, solutions were injected. PBS served as vehicle control.

### Measurement of orofacial hypersensitivity

The mechanical hypersensitivity of the orofacial region was assessed as previously described ^19^. The mice were habituated to small cages for 3 days, then habituated to an area on a table for an additional 3 days. During the last 3 days, mice were also habituated to von Frey filament (0.07-0.6g) stimulations. Naive mice that did not respond to 0.6g von Frey filaments, which is considered baseline, were selected for the experimental procedure ^50^. Experimental mice were probed with 0.008-0.6g filaments to the skin above MM using an up-down approach ^19^. Intervals between von Frey filament applications were varied but >30 sec. Mechanical thresholds were assessed and calculated as previously described ^19, 50^.

### RNA seq transcriptomic data generation, analyses, and statistics

Fresh MM tissues were homogenized in Rn-easy solution (Qiagen) using a Bead Mill Homogenizer (Omni International, Kennesaw, GA). Extraction of RNA, RNA quality and integrity control, cDNA library preparation with oligo-dT primers following the Illumina TruSeq stranded mRNA sample preparation guide (Illumina, San Diego, CA), and sequencing procedure with Illumina HiSeq 3000 platform with 30-50x10^6^ bp reading depth were previously described in detail ^19, 20^. Post-sequencing de-multiplexing with CASAVA, the combined raw reads were aligned to mouse genome build mm9/UCSC hg19 using TopHat2 default settings and differentially expressed genes (DEGs) were identified using DESeq2 after performing median normalization, also previously described in detail ^19, 20^.

Quality control statistical analysis of outliers, intergroup variability, distribution levels, PCA, and hierarchical clustering analysis were performed to statistically validate the experimental data. Multiple correction test was performed with the Benjamini-Hochberg procedure and adjusted p-value (Padj) was generated. If not specified in the text, criteria for selecting DEGs were expression levels with RPKM>1, fold-change (FC)>2, and statistically significant DEGs with Padj<0.05. Venn diagrams were generated using https://bioinfogp.cnb.csic.es/tools/venny/. Genes were clustered according to biological processes using the PANTHER software (http://www.pantherdb.org/).

### Flow cytometry

Flow cytometry was used to assess immune cell profiles in MM biopsies. To eliminate the contributions of immune cells from blood, mice were perfused with cold PBS prior to tissue dissections. Single-cell suspensions from MM biopsies were generated by treating tissues for 70 min at 37°C with 250 μg/ml Liberase (Millipore-Sigma) and 100 μg/ml dispase II (Millipore-Sigma), washing with DMEM media containing 5% FCS, triturating with Pasteur pipettes, and then filtering through a 70μm strainer. Single-cell suspensions were resuspended and centrifuged in isotonic 40% Percoll (Cytiva, Sweden) to enrich live immune cells. Cell suspensions were first stained for viability using Zombie NIR™ Fixable Viability Kit (BioLegend, San Diego, CA) for 20 min at room temperature in PBS combined with FcR blocking antibody (1 μg, clone 2.4G2, BioXCell, Missouri, TX) to block non-specific binding. Cells then were washed with 2% FBS/PBS and stained with antibodies against surface antigens for 30 min on ice. Fluorochrome-conjugated antibodies against mouse CD45 (clone 30-F11), CD3 (145-2C11), B220 (RA3-6B2), CD11b (M1/70), CD64 (X54-5/7.1), CD11c (N418), MHC-II (M5/114.15.2), Ly-6G (1A8), Ly-6C (KH1.4) were purchased from BioLegend (San Diego, CA), eBioscience (San Diego, CA) or BD Biosciences (San Jose, CA). Flow cytometry was performed using Celesta or LSRII cytometer (BD Biosciences; San Jose, CA). Data was analyzed using FlowJo LLC v10.6.1 software. The gating strategy used to select immune populations in MM was done as previously described ^51^. Live CD45^+^ cells were gated using the markers listed below to define cell populations: neutrophils (Nph, CD11b^+^/Ly6G^+^); macrophages (Mph, CD11b^+^/MHCII^hi^/CD64^+^); inflammatory Mph (iMph, CD11b^+^/MHCII^hi^/CD64^+^/Ly6C^+^); monocytes (Mo, CD11b^+^/MHCII^lo^/SSC^lo^/CD64^+^); inflammatory Mo (iMo, CD11b^+^/MHCII^lo^/SSC^lo^/CD64^+^/ Ly6C^hi^); B cells (B, B220^+^/CD11b^-^/CD11c^-^); T cells (T, CD3^+^/CD11b^-^/CD11c^-^), NK (Natural killer cells NK1.1^+^TCRb^-^), CD11b^+^ DCs (Dendritic cells; CD11b^+^CD64^-^CD24^hi^MHCII^hi^CD11c^+^); and CD11b^-^ DCs (CD11b^-^CD64^-^CD24^hi^MHCII^hi^CD11c^+^).

### Immunohistochemistry (IHC)

MM were dissected from 4% paraformaldehyde perfused Ccr2^RFP^/Cx3cr1^GFP^ mice, fixed additionally with 4% paraformaldehyde for 20 min, cryoprotected with 10% and then 30% sucrose in phosphate buffer and then embedded in Neg-50^TM^ (Richard Allan Scientific, Kalamazoo, MI). Cryo-sections of MM (30-35 μm) were collected for IHC. IHC was carried out as previously described ^19, 52^. The following antibodies were used in the study: anti-rat monoclonal S100A8 (Cat: MAB3059; R&D Systems; Minneapolis, MN) and anti-goat Iba1 (Cat: NB100-1028; Novus; Centennial, CO). For non-conjugated primary antibodies, sections were incubated with species-appropriate donkey Alexa Fluor secondary antibodies (1:200; Jackson Immuno-Research, West Grove, PA). Control IHC was performed on tissue sections processed as described but either lacking primary antibodies or lacking primary and secondary antibodies.

Z-stack IHC images were acquired from 3-5 independent tissue sections from 3 animals using a Keyence BZ-X810 All-in-One Fluorescent Microscope (Keyence, Itasca, IL) or a Nikon Eclipse 90i microscope (Melville, NY, USA) equipped with a C1si laser scanning confocal imaging system. Images were processed with NIS-elements software (Nikon Instruments, Melville, NY) or Adobe Photoshop CS2 software. The scale of microphotographs were placed by NIS-elements or Keyence software.

## Results

### CFA-induced transcriptomic changes in the MM

One of the aims of this study was to define whether Col-induced transcriptomic changes in MM are distinct compared to CFA treatment. Thus, our studies began with CFA MM injections. At 1d post-CFA, mice displayed mechanical hypersensitivity of the orofacial area compared to baseline, which was assessed as previously described (1-way ANOVA; F (2, 21) = 39.12; P<0.0001; n=8; *Suppl Fig 1A*)^19^. We then isolated MM1d post CFA (n=3) and performed bulk RNA-seq as previously described^20^. Considering selections as RPKM>1, FC>2 and Padj<0.05, 196 differentially expressing genes (DEGs) were up-regulated and 78 down-regulated at 1d post CFA treatment. As expected, CFA triggered extensive pro-inflammatory processes in MM at day 1 post injected (*Fig 1A*). In contrast, CFA treatment down-regulated DEGs did not fit into biological process according to statistical over-representation test. Among up-regulated DEGs, there are many cytokines, chemokines and their receptors, receptors, such as *Il6, Tnf, Ccl2, Ccr2, Il1b* known to be involved in sensitizing sensory neurons and/or the nociceptive pathway (*Table 1*). Other notable up-regulated DEGs related to the immune system were *Tlr13, Tlr4, Tlr1, Il33, Csf3, Csf2rb, Fcgr1, Nfkbid, Cd300lf, Selp, Oas2, Irf5, Il2rg*, and *Runx1*. Overall, up-regulated DEGs point toward acute inflammatory processes (*Saa1, Saa3, Tlr4, Tlr1*), chemotactic activity for monocytes (*Ccl2, Ccr2, Ccl7*), inflammatory monocytes (*Ccl3, Ccl4, Ccl8, Ccl12*), neutrophils (*Cxcl2, Cxcl3, Cxcl5, Ccl9*), and low levels of phagocytic activity (*Clec4e, Msr1*). Data also revealed that there were only few DEGs related to macrophage and natural killer cell chemotactic pathways.

**Figure 1:**
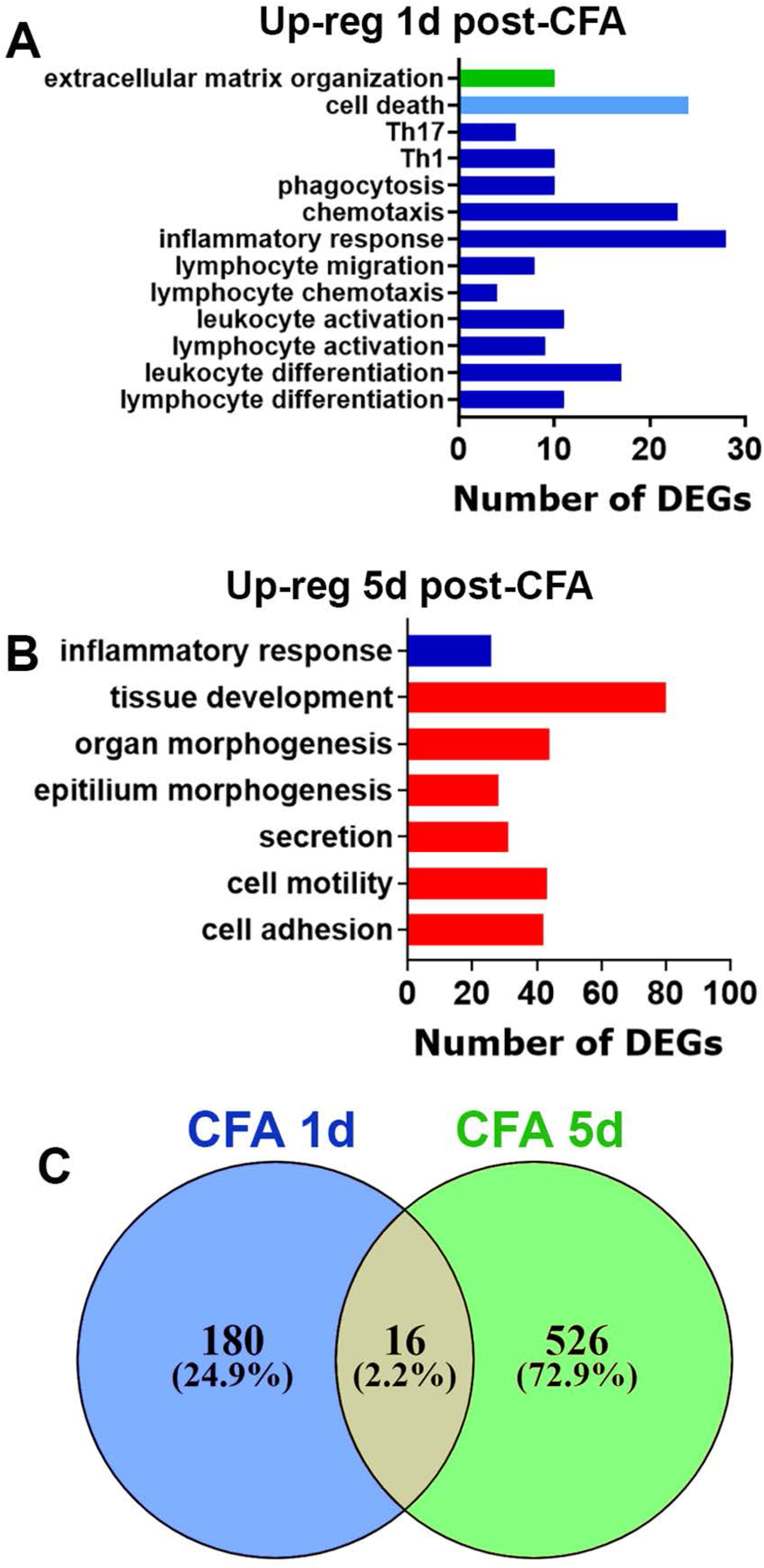
CFA-induced regulation of differential expressed genes (DEGs) in the masseter muscle (MM). (**A**) Biological processes for up-regulated DEGs at 1d post-CFA. (**B**) Biological processes for up-regulated DEGs at 5d post-CFA. (**C**) Venn diagrams of CFA-induced up-regulation of DEGs in the MM at 1d and 5d post-CFA. X-axis on *the panels A, B* represents numbers of DEGs. Y-axis notes biological processes.

**Table 1.** Examples of up-regulated DEGs in MM at 1d post-CFA injection.

| **ID** | **BL** | **CFA 1d** | **FC** | **Name** |
| --- | --- | --- | --- | --- |
| *Cxcl3* | 0 | 80.00 | Inf | chemokine (C-X-C motif) ligand 3 |
| *Cxcl2* | 0.3 | 382.5 | 1062.4 | chemokine (C-X-C motif) ligand 2 |
| *Saa3* | 1.9 | 1438.1 | 733.9 | serum amyloid A 3 |
| *Ccl3* | 0.9 | 102.2 | 107.3 | chemokine (C-C motif) ligand 3 |
| *Saa1* | 4.0 | 315.2 | 77.1 | serum amyloid A 1 |
| *Ccl7* | 8.4 | 604.9 | 71.3 | chemokine (C-C motif) ligand 7 |
| *Cxcl5* | 2.1 | 148.3 | 69.4 | chemokine (C-X-C motif) ligand 5 |
| *Ccl4* | 2.2 | 102.3 | 44.8 | chemokine (C-C motif) ligand 4 |
| *Il6* | 1.2 | 55.3 | 44.0 | interleukin 6 |
| *Ccl12* | 3.1 | 99.5 | 31.8 | chemokine (C-C motif) ligand 12 |
| *Il1b* | 19.0 | 557.5 | 29.3 | interleukin 1 beta |
| *Ccl2* | 18.4 | 398.2 | 21.6 | chemokine (C-C motif) ligand 2 |
| *Clec4n* | 6.2 | 134.4 | 21.3 | C-type lectin domain family 4, member n |
| *Clec4d* | 14.3 | 305.3 | 21.2 | C-type lectin domain family 4, member d |
| *Arg1* | 39.5 | 658.9 | 16.6 | arginase |
| *Osm* | 4.7 | 74.9 | 15.7 | oncostatin M |
| *Msr1* | 29.0 | 296.7 | 10.2 | macrophage scavenger receptor 1 |
| *Tnf* | 5.3 | 51.1 | 9.6 | tumor necrosis factor |
| *Ccl8* | 33.6 | 295.0 | 8.8 | chemokine (C-C motif) ligand 8 |
| *Ccr5* | 32.5 | 262.7 | 8.1 | chemokine (C-C motif) receptor 5 |
| *Ccr1* | 47.3 | 301.3 | 6.4 | chemokine (C-C motif) receptor 1 |
| *Aif1* | 21.1 | 134.0 | 6.3 | allograft inflammatory factor 1 |
| *Ccl9* | 329.0 | 1840.0 | 5.6 | chemokine (C-C motif) ligand 9 |
| *Ccr2* | 182.0 | 712.9 | 3.9 | chemokine (C-C motif) receptor 2 |

At 5d post-CFA, mechanical response thresholds returned to baseline levels (*Suppl Fig 1A*). MM was isolated at 5d post CFA (n=3) and bulk RNA-seq was performed^20^. Using the same selection criteria as above, 542 DEGs were up-regulated and only 18 down-regulated, which did not fit into any biological process (*Fig 1C*). Inflammatory processes were substantially resolved. However, a set of DEGs (*Saa3, Ccl3, Aif1*) up-regulated at 1d post-CFA were still present. Additionally, distinct immune system related DEGs compared to 1d post-CFA (*Cxcl17, Cx3cr1, Tlr3, Gzma, Itgax*) were upregulated at 5d post-CFA (*Fig 1B*). Generally, as low as 16 upregulated DEGs overlapped when 1d compared to 5d post-CFA time points (*Fig 1C*). Biological processes at 5d post-CFA, when mechanical responses were on baseline levels, were dominated by genes involved in various cellular and extracellular matrix repair and developmental processes (*Fig 1B*). In summary, biological processes at 1d (mechanical hypersensitivity) and 5d post-CFA (return to baseline nociception level) were found to be significantly different. Transcriptomic changes in the MM triggered at 1d post-injection of CFA were mainly represented by genes controlling chemotactic activity for monocytes and neutrophils. DEGs were linked to tissue repair processes and only few pro-inflammatory genes were up regulated in MM at 5d post administration.

### Low dose (0.2U) collagenase-induced transcriptomic changes in the MM

Tissue damage was mimicked with a low (0.2U) or high (10U) dose MM injections of Col. Low (0.2U) dose of Col injections induced mechanical hypersensitivity in orofacial area, which was detected at 1d, but not 5d post injection (1-way ANOVA; F (2, 21) = 23.05; P<0.0001; n=8; *Suppl Fig 1B*). We then isolated MM at 1d and 5d post Col (n=3-4) and bulk RNA-seq was performed as previously described^20^. For 1d post-Col and RPKM>1, FC>2 and Padj<0.05 selection, 93 DEGs were up-regulated and 11 down-regulated (*Fig 2C*). Up-regulated DEGs upon 0.2U Col treatment showed biological processes associated with tissue repair, including extracellular matrix reorganization and epithelial cell development (*Fig 2A*). Many of the upregulated DEGs were related to epidermis development (*Krt14, Ktr15, Krt17, Krt77, Ktr79, Fgfr3, Evpl*) or extracellular matrix organization (*Col9a1, Col9a2, Col9a3, Col10a1, Wnt3a, Mmp20*). Interestingly, low Col dosage did not trigger upregulation of inflammatory processes at 1d post-administration (*Fig 2A*). Down regulated DEGs by 0.2U Col did not highlight any biological process according to a statistical over-representation test. At 5d post-Col, only 18 DEGs were up-regulated and 8 down-regulated. It seemed that the repair process was completed by 5d post-Col (0.2U) (*Fig 2B*). Only two DEGs remained up-regulated (*Ucp1, Elov16*) and only two down-regulated (*Slc2a3, Hs3st5*) after 5d post-Col (*Supplementary material*). Overall, low dosage of Col (0.2U) produced acute orofacial hypersensitivity which was linked to tissue repair, but not inflammatory processes. These results suggest that tissue restoration was fully completed 5 days post injection.

**Figure 2:**
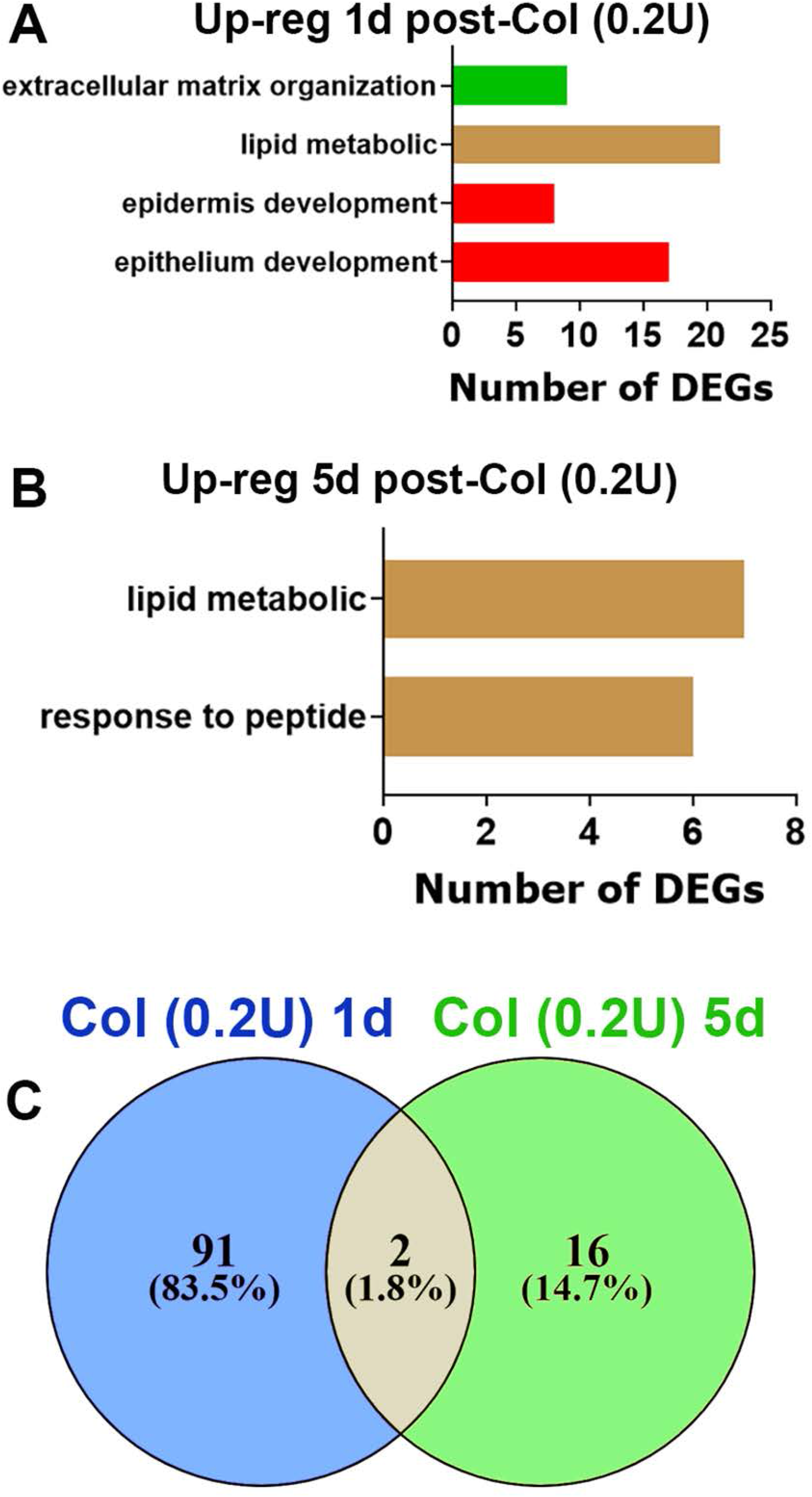
Collagenase type 2 (Col; 0.2U)-induced regulation of DEGs in the MM. (**A**) Biological processes for up-regulated DEGs at 1d post-Col (0.2U). (**B**) Biological processes for up-regulated DEGs at 5d post-Col (0.2U). (**C**) Venn diagrams of Col (0.2U)-induced up-regulation of DEGs in the MM at 1d and 5d post-Col (0.2U). X-axis on *the panels A, B* represents numbers of DEGs. Y-axis notes biological processes.

### High dose (10U) collagenase-induced transcriptomic changes in the MM

At a high dose of Col (10U) we found that mechanical hypersensitivity lasted at least 2 weeks with day 6 post-injection as a bifurcation point in the hypersensitivity developmental trajectory (1-way ANOVA; F (3, 28) = 68.00; n=8; P<0.0001; *Suppl Fig 1B*). Accordingly, we isolated MM (n=3-4) at 1d, 6d and 14d post Col (10U) and evaluated transcriptomic at these time points. DEGs selection criteria was as above. At 1d post-Col, 334 DEGs were up and 464 DEGs were down-regulated. Inflammatory processes dominated up-regulated DEGs at 1d post-Col (*Fig 3A*). Down-regulated DEGs were associated with muscle damage and reductions of aerobic cellular respiration (*Fig 3B*). At 6d post-Col, as many as 1623 DEGs were up, and 1648 DEGs down-regulated. Analysis showed that at 6d post-Col, which is the last time point when hypersensitivity was detected, inflammation was elevated compared to 1d post-injection, tissue repair had begun and lipid metabolism was reduced, while cellular respiration and metabolism had yet to be recovered (*Figs 3C, 3D*). Upon orofacial myogenous mechanical hypersensitivity resolution at 14d post-Col, only 99 DEGs were up- and 65 down-regulated. It appeared that biological processes returned to homeostasis and none of them was up or down-regulated.

**Figure 3:**
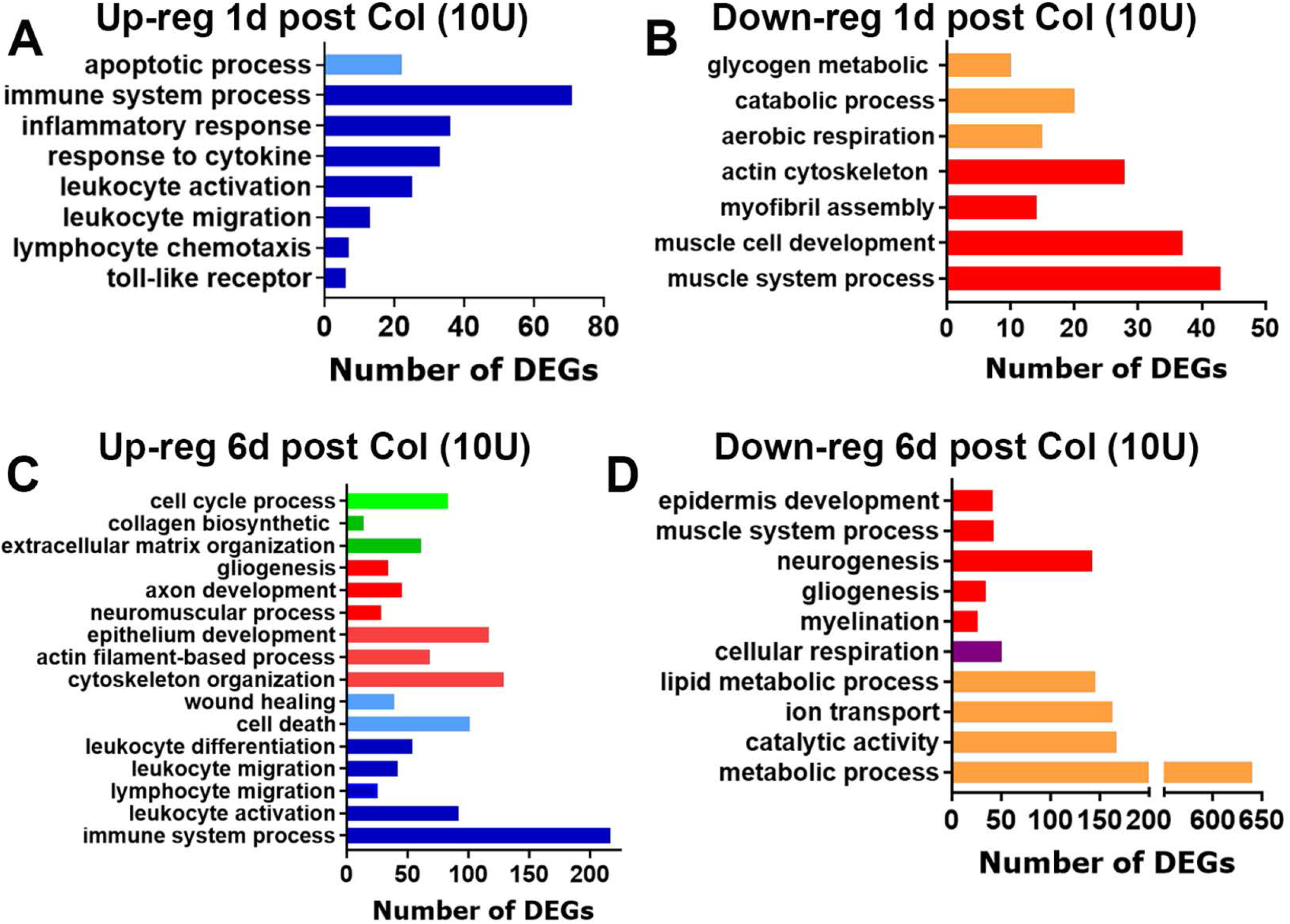
Up- and down-regulated biological processes in the MM after 10U Col treatments. (**A**) Biological processes for up-regulated DEGs at 1d post-Col (10U). (**B**) Biological processes for down-regulated DEGs at 1d post-Col (10U). (**C**) Biological processes for up-regulated DEGs at 6d post-Col (10U). (**D**) Biological processes for down-regulated DEGs at 6d post-Col (10U). The X-axis on *the panels A-D* represents numbers of DEGs. The Y-axis notes biological processes.

Next, we analyzed differences in inflammatory mediators (cytokines and chemokines) and processes at 1d post-injection of Col compared to CFA. CFA, 0.2U Col, and 10U Col initially (1d post-injection) produced substantially different expression changes in MM (*Figs 4A, 4B*). However, like CFA, 10U Col treatment up-regulated certain numbers of the same pro-inflammatory genes (*Table 2*). Thus, common up-regulated DEGs for CFA and 10U Col intramuscular treatments included monocyte and neutrophil associated genes: *Cxcl2, Cxcl3, Cxcl5, Ccl2, Ccl7, Ccl8, Ccl12, Ccr5, Ccr2, Aif1, Mrc1, Msr1, Ms4a6b, Clec4d, Clec4n, Fcgr1, Mt2, Mcoln2, Runx1, Tlr1, Tlr4*, etc. One of the main differences in biological processes at 1d post-injection between CFA and 10U Col treatment was that, unlike CFA, 10U Col administration led to up-regulation of genes linked to macrophage activation and the down-regulation of DEGs associated with muscle structure, muscle cell development and catabolic processes (*Fig 3B*).

**Figure 4:**
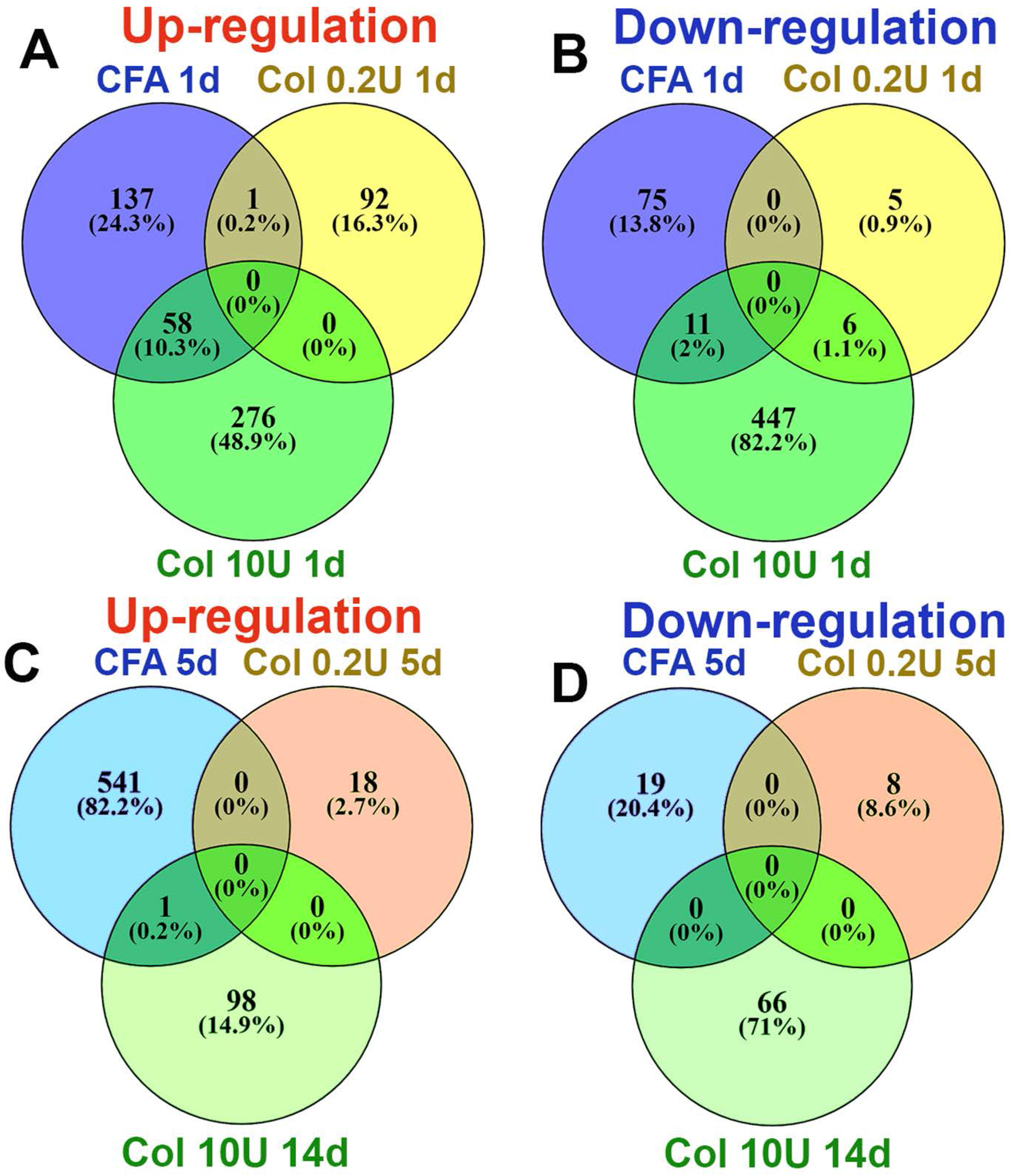
Comparisons of up- and down-regulated DEGs in the MM after CFA, 0.2U Col or 10U Col treatments. (**A**) Venn diagrams of CFA, 0.2U Col and 10U Col-induced up-regulation of DEGs in the MM at 1d post-treatments. (**B**) Venn diagrams of CFA, 0.2U Col and 10U Col-induced down-regulation of DEGs in the MM at 1d post-treatments as indicated on panels. (**C**) Venn diagrams of CFA, 0.2U Col and 10U Col-induced up-regulation of DEGs in the MM at 5d post-treatments. (**D**) Venn diagrams of CFA, 0.2U Col and 10U Col-induced down-regulation of DEGs in the MM at 5d post-treatments as indicated on panels.

**Table 2.**
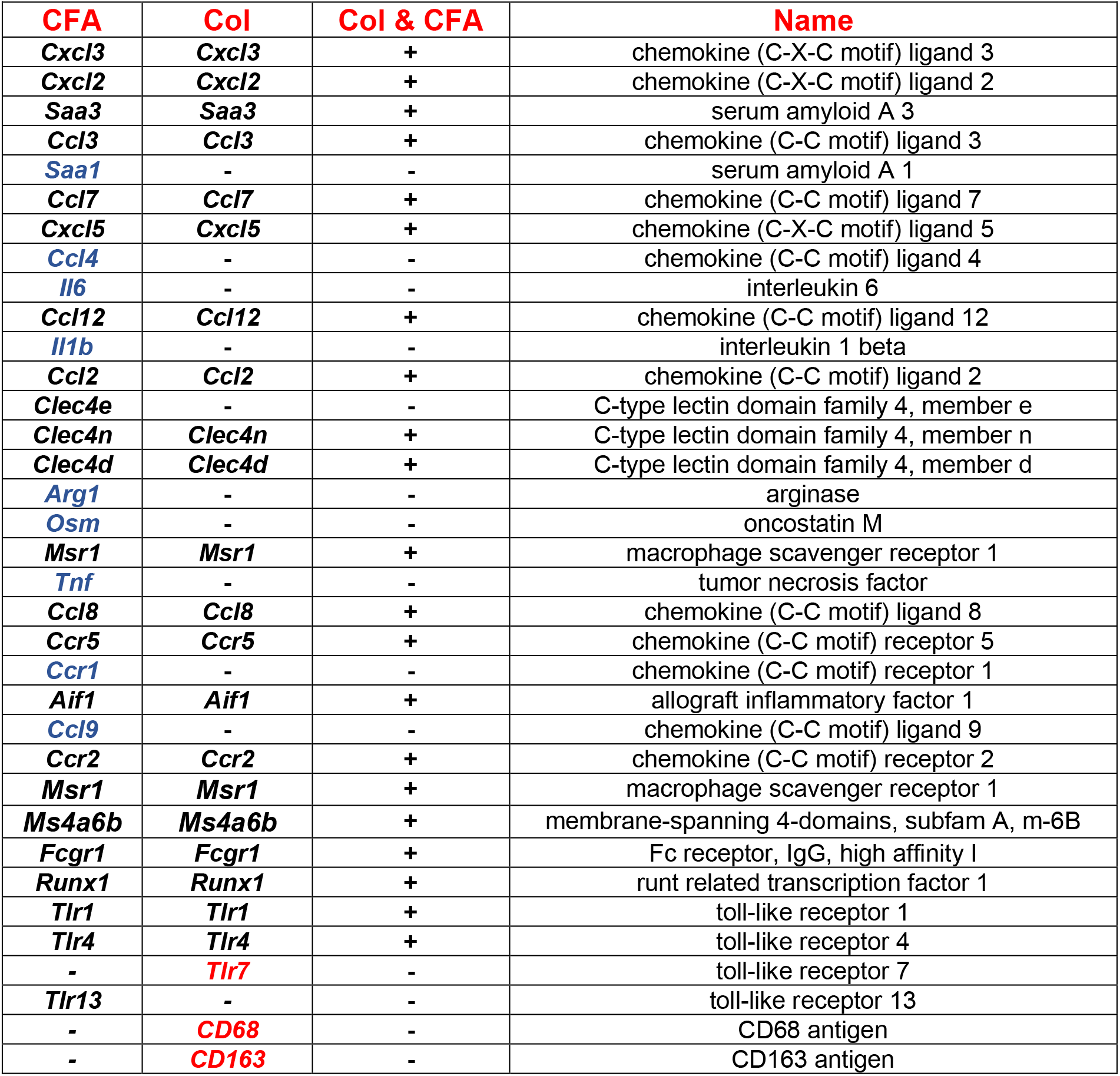
Examples of up-regulated DEGs in MM at 1d post-CFA or Col (10U) injection.

| **CFA** | **Col** | **Col & CFA** | **Name** |
| --- | --- | --- | --- |
| *Cxcl3* | *Cxcl3* | + | chemokine (C-X-C motif) ligand 3 |
| *Cxcl2* | *Cxcl2* | + | chemokine (C-X-C motif) ligand 2 |
| *Saa3* | *Saa3* | + | serum amyloid A 3 |
| *Ccl3* | *Ccl3* | + | chemokine (C-C motif) ligand 3 |
| *Saa1* | - | - | serum amyloid A 1 |
| *Ccl7* | *Ccl7* | + | chemokine (C-C motif) ligand 7 |
| *Cxcl5* | *Cxcl5* | + | chemokine (C-X-C motif) ligand 5 |
| *Ccl4* | - | - | chemokine (C-C motif) ligand 4 |
| *Il6* | - | - | interleukin 6 |
| *Ccl12* | *Ccl12* | + | chemokine (C-C motif) ligand 12 |
| *Il1b* | - | - | interleukin 1 beta |
| *Ccl2* | *Ccl2* | + | chemokine (C-C motif) ligand 2 |
| *Clec4e* | - | - | C-type lectin domain family 4, member e |
| *Clec4n* | *Clec4n* | + | C-type lectin domain family 4, member n |
| *Clec4d* | *Clec4d* | + | C-type lectin domain family 4, member d |
| *Arg1* | - | - | arginase |
| *Osm* | - | - | oncostatin M |
| *Msr1* | *Msr1* | + | macrophage scavenger receptor 1 |
| *Tnf* | - | - | tumor necrosis factor |
| *Ccl8* | *Ccl8* | + | chemokine (C-C motif) ligand 8 |
| *Ccr5* | *Ccr5* | + | chemokine (C-C motif) receptor 5 |
| *Ccr1* | - | - | chemokine (C-C motif) receptor 1 |
| *Aif1* | *Aif1* | + | allograft inflammatory factor 1 |
| *Ccl9* | - | - | chemokine (C-C motif) ligand 9 |
| *Ccr2* | *Ccr2* | + | chemokine (C-C motif) receptor 2 |
| *Msr1* | *Msr1* | + | macrophage scavenger receptor 1 |
| *Ms4a6b* | *Ms4a6b* | + | membrane-spanning 4-domains, subfam A, m-6B |
| *Fcgr1* | *Fcgr1* | + | Fc receptor, IgG, high affinity I |
| *Runx1* | *Runx1* | + | runt related transcription factor 1 |
| *Tlr1* | *Tlr1* | + | toll-like receptor 1 |
| *Tlr4* | *Tlr4* | + | toll-like receptor 4 |
| - | *Tlr7* | - | toll-like receptor 7 |
| *Tlr13* | - | - | toll-like receptor 13 |
| - | *CD68* | - | CD68 antigen |
| - | *CD163* | - | CD163 antigen |

At the time point preceding myalgia resolution (6d for 10U Col; *Suppl Fig 1B*), expression and the number of pro-inflammatory DEGs were significantly increased compared to 1d post 10U Col (*Figs 3C, 3D*). The rise in pro-inflammatory DEG numbers accompanied wound healing and tissue repair processes, included partial muscle, glia, neuro-muscular junction, extracellular matrix, and epithelium repair (*Fig 3C*), as well as drastic downregulation of metabolic processes (*Fig 3D*). Overall, expression changes in MM from 1d to 6d post-Col were dramatic (Figs 5A, 5B). Next, we looked for a subset of immune system related DEGs that could be upregulated at 6d compared to 1d post-Col (10U). Thus, at 6d post-Col, biological processes in MM included accumulation of macrophages (*Cx3cr1, Ccl22, mpeg1, lyz2, csfr1*), natural killer (NK; *granzymes, CD53, Cd244*) and natural killer T cells (NKT; *Cxcl16*), dendritic cells (*itgax, Cd48, Cd80, Cd86*) and T-cells (*Cd4, Cd40, Cd72*) (*Table 3*).

**Figure 5:**
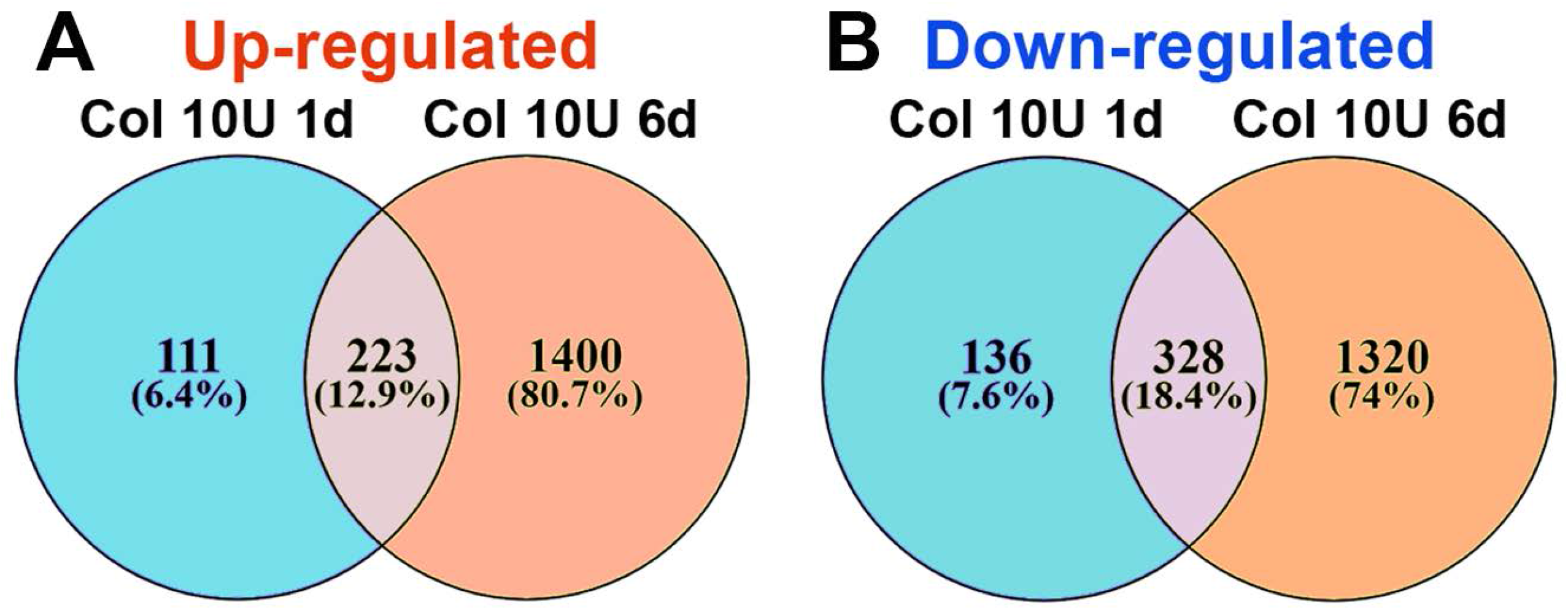
Comparisons of up- and down-regulated DEGs in the MM at different time points after 10U Col treatments. (A) Venn diagrams of 10U Col-induced up-regulation of DEGs in the MM at 1d and 6d post-treatments. (B) Venn diagrams of 10U Col-induced down-regulation of DEGs in the MM at 1d and 6d post-treatments as indicated on panels.

**Table 3.**
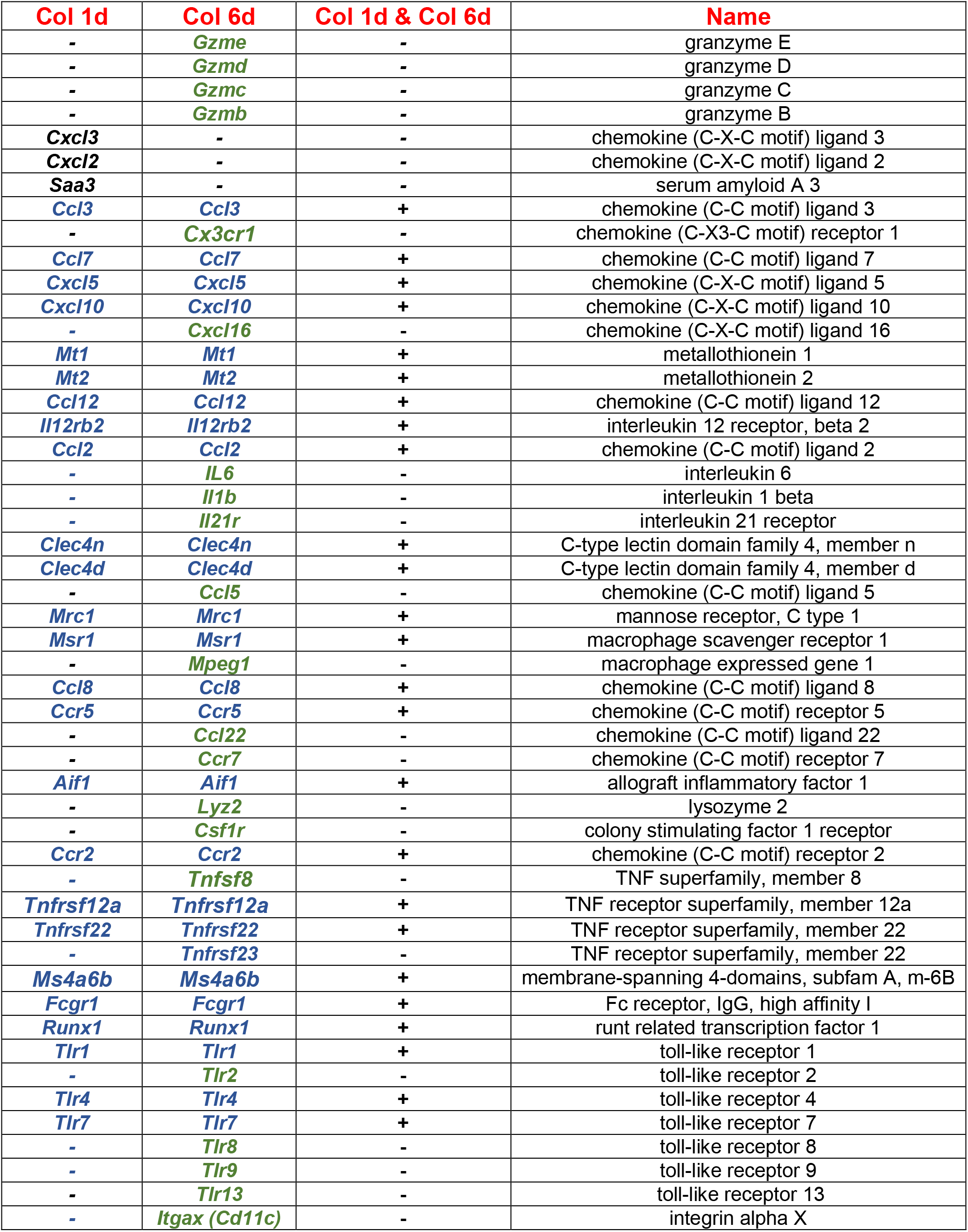
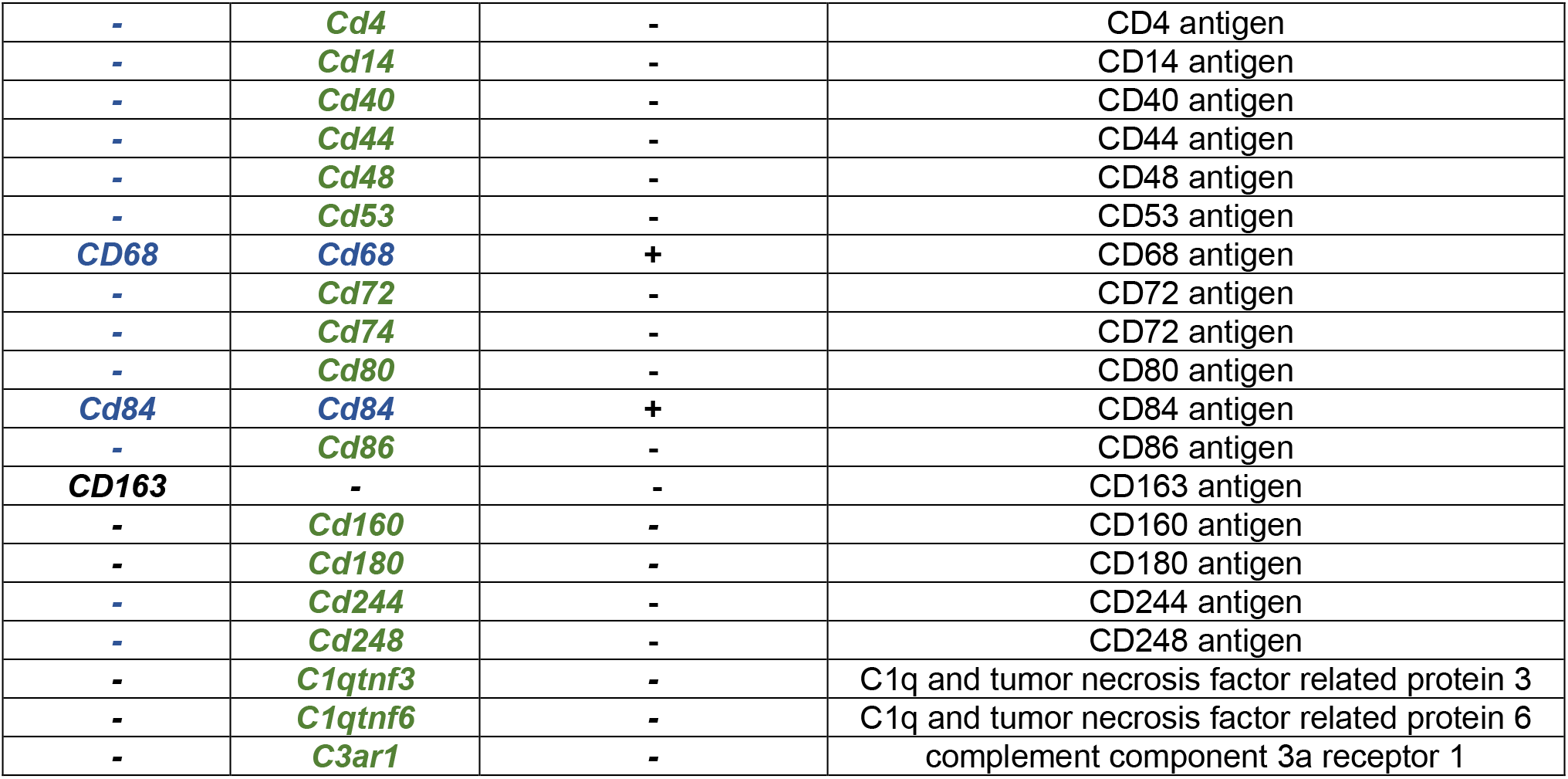
Examples of up-regulated DEGs in MM at 6d compared to 1d post-Col (10U).

| Col 1d | Col 6d | Col 1d & Col 6d | Name |
| --- | --- | --- | --- |
| - | *Gzme* | - | granzyme E |
| - | *Gzmd* | - | granzyme D |
| - | *Gzmc* | - | granzyme C |
| - | *Gzmb* | - | granzyme B |
| *Cxcl3* | - | - | chemokine (C-X-C motif) ligand 3 |
| *Cxcl2* | - | - | chemokine (C-X-C motif) ligand 2 |
| *Saa3* | - | - | serum amyloid A 3 |
| *Ccl3* | *Ccl3* | + | chemokine (C-C motif) ligand 3 |
| - | *Cx3cr1* | - | chemokine (C-X3-C motif) receptor 1 |
| *Ccl7* | *Ccl7* | + | chemokine (C-C motif) ligand 7 |
| *Cxcl5* | *Cxcl5* | + | chemokine (C-X-C motif) ligand 5 |
| *Cxcl10* | *Cxcl10* | + | chemokine (C-X-C motif) ligand 10 |
| - | *Cxcl16* | - | chemokine (C-X-C motif) ligand 16 |
| *Mt1* | *Mt1* | + | metallothionein 1 |
| *Mt2* | *Mt2* | + | metallothionein 2 |
| *Ccl12* | *Ccl12* | + | chemokine (C-C motif) ligand 12 |
| *Il12rb2* | *Il12rb2* | + | interleukin 12 receptor, beta 2 |
| *Ccl2* | *Ccl2* | + | chemokine (C-C motif) ligand 2 |
| - | *IL6* | - | interleukin 6 |
| - | *Il1b* | - | interleukin 1 beta |
| - | *Il21r* | - | interleukin 21 receptor |
| *Clec4n* | *Clec4n* | + | C-type lectin domain family 4, member n |
| *Clec4d* | *Clec4d* | + | C-type lectin domain family 4, member d |
| - | *Ccl5* | - | chemokine (C-C motif) ligand 5 |
| *Mrc1* | *Mrc1* | + | mannose receptor, C type 1 |
| *Msr1* | *Msr1* | + | macrophage scavenger receptor 1 |
| - | *Mpeg1* | - | macrophage expressed gene 1 |
| *Ccl8* | *Ccl8* | + | chemokine (C-C motif) ligand 8 |
| *Ccr5* | *Ccr5* | + | chemokine (C-C motif) receptor 5 |
| - | *Ccl22* | - | chemokine (C-C motif) ligand 22 |
| - | *Ccr7* | - | chemokine (C-C motif) receptor 7 |
| *Aif1* | *Aif1* | + | allograft inflammatory factor 1 |
| - | *Lyz2* | - | lysozyme 2 |
| - | *Csf1r* | - | colony stimulating factor 1 receptor |
| *Ccr2* | *Ccr2* | + | chemokine (C-C motif) receptor 2 |
| - | *Tnfsf8* | - | TNF superfamily, member 8 |
| *Tnfrsf12a* | *Tnfrsf12a* | + | TNF receptor superfamily, member 12a |
| *Tnfrsf22* | *Tnfrsf22* | + | TNF receptor superfamily, member 22 |
| - | *Tnfrsf23* | - | TNF receptor superfamily, member 22 |
| *Ms4a6b* | *Ms4a6b* | + | membrane-spanning 4-domains, subfam A, m-6B |
| *Fcgr1* | *Fcgr1* | + | Fc receptor, IgG, high affinity I |
| *Runx1* | *Runx1* | + | runt related transcription factor 1 |
| *Tlr1* | *Tlr1* | + | toll-like receptor 1 |
| - | *Tlr2* | - | toll-like receptor 2 |
| *Tlr4* | *Tlr4* | + | toll-like receptor 4 |
| *Tlr7* | *Tlr7* | + | toll-like receptor 7 |
| - | *Tlr8* | - | toll-like receptor 8 |
| - | *Tlr9* | - | toll-like receptor 9 |
| - | *Tlr13* | - | toll-like receptor 13 |
| - | *Itgax (Cd11c)* | - | integrin alpha X |
| - | *Cd4* | - | CD4 antigen |
| - | *Cd14* | - | CD14 antigen |
| - | *Cd40* | - | CD40 antigen |
| - | *Cd44* | - | CD44 antigen |
| - | *Cd48* | - | CD48 antigen |
| - | *Cd53* | - | CD53 antigen |
| *CD68* | *Cd68* | + | CD68 antigen |
| - | *Cd72* | - | CD72 antigen |
| - | *Cd74* | - | CD72 antigen |
| - | *Cd80* | - | CD80 antigen |
| *Cd84* | *Cd84* | + | CD84 antigen |
| - | *Cd86* | - | CD86 antigen |
| *CD163* | - | - | CD163 antigen |
| - | *Cd160* | - | CD160 antigen |
| - | *Cd180* | - | CD180 antigen |
| - | *Cd244* | - | CD244 antigen |
| - | *Cd248* | - | CD248 antigen |
| - | *C1qtnf3* | - | C1q and tumor necrosis factor related protein 3 |
| - | *C1qtnf6* | - | C1q and tumor necrosis factor related protein 6 |
| - | *C3ar1* | - | complement component 3a receptor 1 |

We next evaluated gene changes at 14d post-10U Col versus 5d post-0.2U Col. At these time points, hypersensitivity returned to baseline for both treatments (*Suppl Fig 1*). Few DEGs were up- and down-regulated at these time points. Among these few regulated DEGs, almost no similarity could have been detected (*Figs 4C, 4D*). In summary, CFA and Col treatment induced substantially different biological processes in the MM. Additionally, the transcriptomic analysis showed that mechanical hypersensitivity resolution correlates with gradual muscle and extracellular matrix repair, which was supplemented by an elevation in immune system gene expression compared to 1d post 10U Col.

### Collagenase-induced alterations in immune cell profiles in masseter muscle (MM)

To validate bulk RNA-seq data, we used flow cytometry to evaluate immune cell (CD45^+^) profiles in MM at different time points after CFA or Col injections (*Suppl Fig 2*). CFA intra MM treatment increased CD45^+^ cell numbers >14-fold at 1d post-CFA (*t*-test; Veh vs CFA; 2266±77 vs 29300±6084, t=4.443, df=4; P=0.011; n=3). This increase in CD45^+^ cells was due to infiltration/proliferation of monocytes (Mo), inflammatory monocytes (iMo), and neutrophils (Neu) in MM, while the number of B-cells were reduced (2-way ANOVA; interaction F (9, 40) = 42.42; P<0.0001; n=3; *Fig 6A, Suppl Fig 2*). Mph, inflammatory Mph (iMph) as well as natural killer (NK), dendritic cells (DC) and T-cells numbers were not significantly changed at 1d post-CFA (*Fig 6A*).

**Figure 6:**
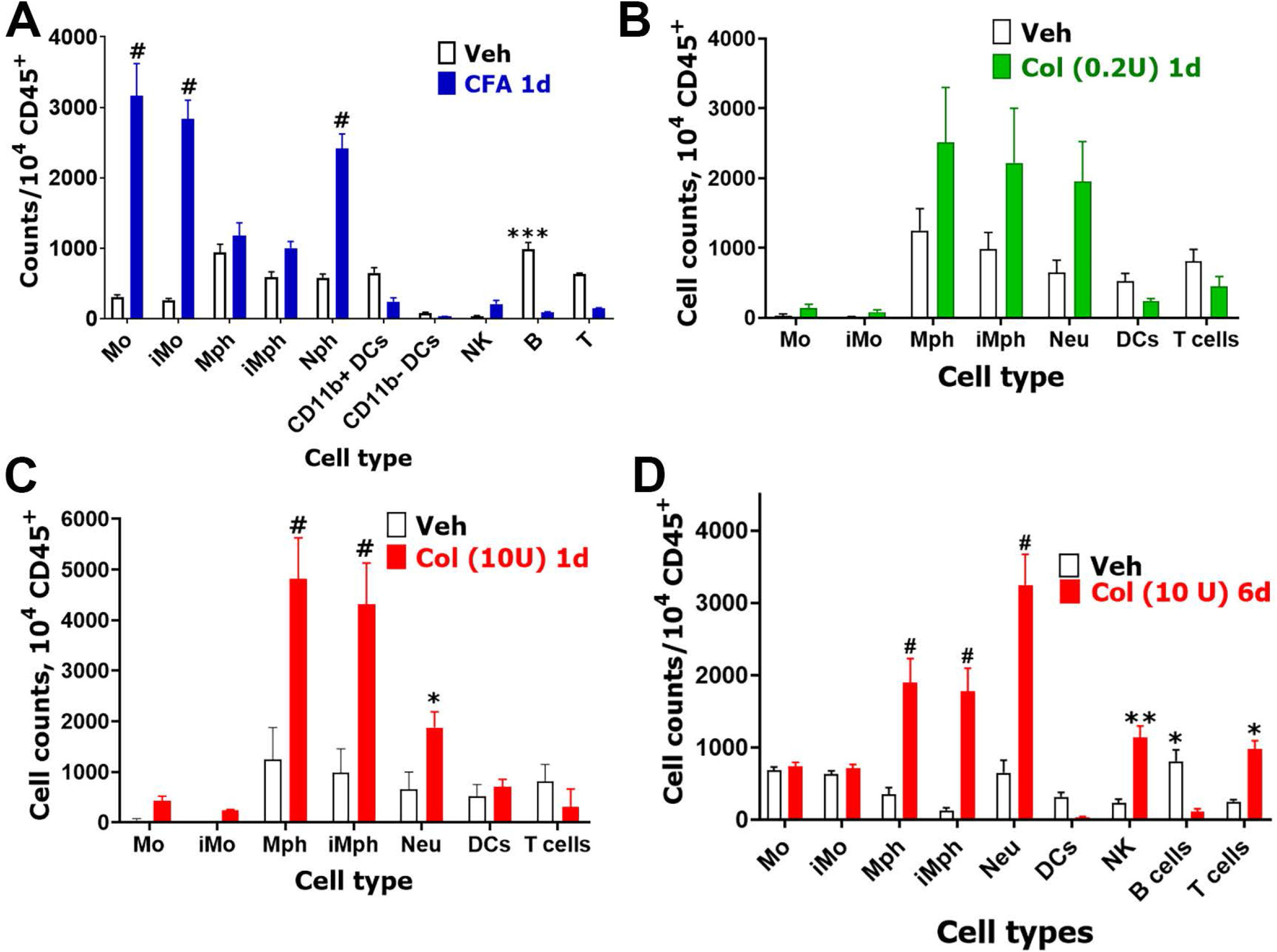
Immune cell profiles in CFA and Col treated MM. **(A)** Immune cell counts per 10^4^ CD45^+^ cells in the MM at 1d post-vehicle (Veh) or CFA single intramuscular treatment. (**B**) Immune cell counts per 10^4^ CD45^+^ cells in MM at 1d post-vehicle (Veh) or Col (0.2U) single intramuscular treatment. (**C**) Immune cell counts per 10^4^ CD45^+^ cells in MM at 1d post-vehicle (Veh) or Col (10U) single intramuscular treatment. (**D**) Immune cell counts per 10^4^ CD45^+^ cells in MM at 6d post-vehicle (Veh) or Col (10U) single intramuscular treatment. DCs – dendritic cells; Mph – macrophages; iMph - inflammatory macrophages; Mo – monocytes; iMo - inflammatory monocytes; NK – natural killer cells; B – B-cells; T – T-cells and Neu – neutrophils. Statistic is 2-way ANOVA (* p<0.05; ** p<0.01; *** p<0.001; # p<0.0001; n=3-6).

Low dose (0.2U) injection of Col slightly increased numbers of CD45^+^ cells, which was statistically significant (*t*-test; Veh vs 0.2U Col; 4965±1040 vs 9928±3671, t=1.301, df=6; P=0.24; n=4, *Fig 6B*). This slight increase was caused by infiltrations/proliferations of macrophages (Mph), inflammatory macrophages (iMph) and Neu in MM, but was statistically insignificant at 1d post-administration (2-way ANOVA; interaction F (6, 42) = 2.24; P=0.058; n=4; *Fig 6B*). An increase in CD45^+^ cells 1d after 10U Col intra-MM injection was substantial, but not as pronounced compared to CFA treatment (*t*-test; Veh vs 10U Col; 2240±777 vs 14980±2648, t=4.105, df=6; P=0.0063; n=4). Moreover, unlike CFA, 10U Col elevated infiltration/proliferation of Mph, iMph and Neu, while Mo and iMo were unaffected in MM (2-way ANOVA; interaction F (6, 42) = 29.70; P<0.0001; n=4; *Fig 6C*).

Since 6d post Col (10U) treatment is a critical time point for the development of chronicity in orofacial hypersensitivity, we performed flow cytometry on MM tissue at 6d-post Col injection and found substantially increased CD45^+^ cell counts (*t*-test; Veh vs 10U Col; 3620±113 vs 9980±337, t=18.87, df=6; P<0.0001; n=4). Besides infiltration of Mph, iMph and Neu at 1d post Col, 6d post-10U Col elevated NK and T cells (2-way ANOVA; interaction F (8, 54) = 19.37; P<0.0001; n=4; *Fig 6D*). These data verify bulk RNA-seq results, which implied elevation of Mph, iMph and Neu as well as NK and T cells at 6d post-Col (10U).

Finally, we performed flow cytometry on MM tissue after resolution of orofacial hypersensitivity for CFA-treatment (resolution found to be at 5d post-injection) and Col-treatment (0.2U or 10U, resolution found to be at 5d and 14d post-administration, respectively) (*Suppl Figs 1A, 1B*). After CFA-induced hypersensitivity resolution, no elevation of CD45^+^ was detected in MM compared to vehicle treatment (*t*-test; Veh vs CFA; 4125±728 vs 6944±564, t=0.3000, df=7; P=0.77; n=3-6). Examination of different immune cell types showed that changes in immune cell profiles at 5d post-CFA compared 5d post vehicle were insignificant (*Suppl Fig 3A*). Similar to CFA, 14d-post Col 0.2U injection led to no increase in CD45^+^ cell counts (*t*-test; Veh vs 0.2U Col; 558±90 vs 657±229, t=0.4623, df=8; P=0.66 n=4-6). As expected, immune cell profiles did not undergo changes (1-way ANOVA; F (6, 56) = 1.254; F (6, 56) = 1.254; P=0.29; n=4-6; *Suppl Fig 3B*). However, MM at 14d post 10U Col injection still had a statistically significant increase of CD45^+^ cell counts (*t*-test; Veh vs 10U Col; 558±90 vs 1450±196, t=4.639, df=8; P= 0.0017; n=4-6). Elevation of CD45^+^ was mainly due to Neu, which remained elevated in MM at 14d post-10U Col injection (2-way ANOVA; interaction F (6, 49) = 8.201; P<0.0001; n=4-6; *Suppl Fig 3B*). In summary, CFA-induced inflammation was principally different compared to Col-triggered inflammation. When compared to a 10U dose of Col, CFA produced a significant elevation of different types of myeloid cells at the initial stage (1d post-injection) (*Fig 6*). Furthermore, the pre-resolution stage (6d) after Col (10U) treatment was characterized by significant activation and infiltration/proliferation of myeloid cells as well as NK and T cells in MM (*Fig 6D*). Finally, at 14d post-Col, when mechanical sensitivity was resolved (*Suppl Fig 1B*), the number of CD45^+^ cells in MM did not return to vehicle-treated levels due to Neu, which remained elevated (*Suppl Fig 3B*).

To further validate our sequencing and flow cytometry data, immunohistochemistry (IHC) was used to define the effect of 10U Col injection into the MM on immune cell populations. We focused on three macrophage/monocyte (Mph/Mo/Neu) markers: CX3CR1, CCR2, Iba1 (aka *Aif1*) and S100a8. Accordingly, vehicle or Col (10U) was injected into MM of Cx3cr1-GFP/Ccr2-RFP reporter mice. One day post-injection MM were dissected and used for IHC staining for Iba1 (marker for Mph) or S100a8 (marker for Neu/Mo) ^21^. CCR2, CX3CR1, and S100a8 were almost absent in the MM of vehicle-injected mice (*Figs 7A, 7A”*; data not shown). Iba1 was detected between muscle fibers in control mice (*Figs 7A, 7A’*). Col injection into the MM triggered the appearance of multiple CCR2^+^, CX3CR1^+^, and S100a8^+^ cells (*Figures 7A, 7A” vs 7B, 7B”*, data not shown), as well as a visible up-regulation of Iba1^+^ cells (*Figs 7A, 7A’ vs 7B, 7B’*). Altogether, these data showed that intra-MM Col injection up-regulated multiple myeloid cells in the MM.

**Figure 7:**
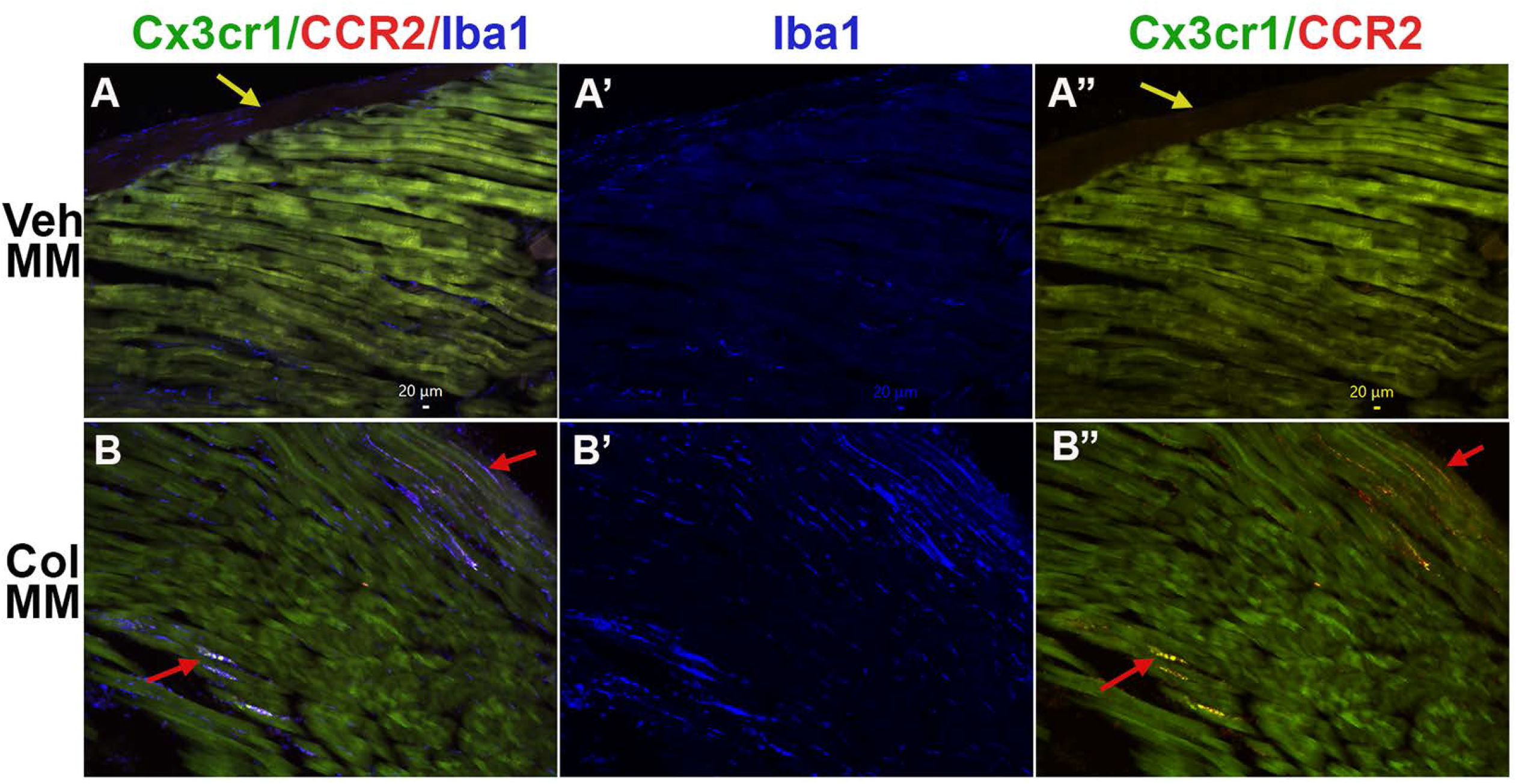
Expression of macrophage/monocyte markers in the MM treated with Col (10U). (**A-B”**) The MM of Cx3cr1-GFP/CCR2-RFP reporter mice intramuscularly treated with Veh (Veh MM; *panels A-A”*) or 10U Col (Col MM; *panels B-B”*) labeled with Iba1. Labeling on each panel is indicated. Yellow arrows on *panels A* and *A”* point to the fascia. Red arrows on the *panels B* and *B”* point to immune cells localized between muscle fibers.

## Discussion

The pathogenesis of myogenous temporomandibular disorder (TMDM) remains largely unknown. However, there is widespread opinion that TMDM is a consequence of tissue damage caused by consistent, repetitive, and prolonged physical overload of the muscles of mastication ^1, 5, 10^. Tissue damage and accompanying muscle ischemia leads to mild inflammation that is characterized by release of inflammatory mediators, such as neuropeptides, serotonin (5-HT), and cytokines that may activate and sensitize nociceptors on peripheral sensory afferents to induce muscle pain and allodynia^22^. Microdialysis sampling mediators in MM showed that IL-1β, GM-CSF, IL-6, IL-7, IL-8 and IL-13 were higher in TMDM patients ^1^. Other small sample sized studies confirmed these findings on significant elevation of cytokines in active trigger points of muscles during myalgia^8, 23^. However, the largest studies found no elevation in cytokines levels during chronic myalgia ^6, 7^. Overall, there is a consensus that chronic and persistent TMDM correlates with mild inflammation. However, there is no clear picture on what type of inflammation and biological processes are present in masticatory muscles. In this regard, we mimicked inflammation two ways: a direct induction with CFA or recreating tissue damage in MM by disrupting the extracellular matrix with collagenase type-2 (Col). Next, we evaluated transcriptional changes in MM at different time points during the development of orofacial hypersensitivity. Furthermore, using flow cytometry and IHC, we examined infiltration/proliferation of immune cells at the same time points after CFA or Col injections. These several approaches allowed us to draw conclusions on the biological processes occurring in MM after CFA or Col injections.

To keep focus of this study, we concentrated on understanding the biological processes in the MM of males, who represent 1/3 TMDM patients. CFA-induced orofacial hypersensitivity was transient and lasted up to 5d. When hypersensitivity resolved at 5d, biological processes related to tissue repair were still present. However, inflammatory processes were mostly receded. In contrast, at the initial stage, (1d post-CFA) hypersensitivity was apparent, and inflammatory processes were dominant in MM. At this stage post-CFA, active immune cells were composed of infiltrating monocytes (Mo), inflammatory monocytes (iMo) and neutrophils (Neu). Importantly, bulk RNA-seq and flow cytometry data indicate that CFA triggers minimal (if any) changes in macrophage activity. A similar pattern in the immune system behavior was noted after CFA injection into the paw ^13^.

Treatment with low dose Col generated weak/low inflammation with slight up-regulation of *Cd68, Ccl2, Ccl7*, but not pro-inflammatory interleukins, such as IL-1β and IL-6. It is likely that 10U Col is more representative of MM damage than 0.2U Col. Moreover, 0.2U Col did not trigger substantial inflammatory processes, while 10U Col injection produced drastic activation and/or infiltration of immune cells. Unlike CFA, dominant cells were macrophages (Mph) and inflammatory macrophages (iMph) alone with Neu at 1d post-Col (10U). Similar behavior of the immune system was observed in the paw after incision surgery^13^. Another key difference of Col (10U) compared to CFA treatments was a significant down-regulation of genes necessary for muscle tissue repair, including *Casq1, Myom2, Myh7, Mylk2, Myoz1, Lmod1, Acta1, Actn3, Smtn, Tpm1, Tcap, Tmod4, Tnnt3, Tuba8*, etc^24–27^.

TMDM is a chronic pain condition^28^. Hypersensitivity after single treatment with 10U Col was eventually resolved by 12-14d. Six days post Col treatment is a critical bifurcation point preceding pain resolution. Hence, understanding the biological processes in MM at this time point is important. Compared to 1d post-Col, at 6d post-Col, up-regulation of DEGs related to tissue repair and elevation of immune system related gene expressions in MM was noted. Up-regulation of *Cx3cr1, Ccl22, mpeg1, lyz2, csfr1* indicated an additional activation and/or accumulation of pro-inflammatory macrophages ^29, 30^. *Cxcl16* is produced by dendritic cells and is chemoattractant for NKT cells^31^. *Granzymes, CD53 and Cd244* up-regulation implies activation/infiltration of NK cells^32, 33^. *Itgax* (aka *CD11c*), as well as *Cd48, Cd80,* and *Cd86* relate to DC^34, 35^. Finally, *Cd4* and *Cd40* are associated with T-cells^36^. Overall, some immune system related genes were present at 1d post-Col, but their expression was dramatically increased at 6d post-Col. These genes are mainly associated with macrophage activation. Moreover, from 1d to 6d post-10U Col, we observed elevation of genes linked to activation/accumulation of NK, NKT and T-cells.

The inflammatory response could play a central role in bridging the initial response to muscle injury to that of timely muscle injury repare^37^ or lead to persistent tissue damage^38, 39^, which contributes to chronic TMDM. Muscle damage induces three pathways that activate inflammatory cells: complement cascade (C3a and C5a) that recruits circulating leukocytes; damage-associated molecular patterns (DAMPs), responsible for recruiting circulating leukocytes to the injured site; and muscle-activated local immune cells releasing cytokines and chemokines^27^. We report here that these processes took place at 1d and/or 6d post-Col (10U) injection into MM (*Table 3*). The cytokines CCL2, CCR2 and some TLRs were present at both time points^40^ (*Table 3*), while CXCL2, IL-1β, C1 and C3 complex components and chemokines, CDs and TLR were encountered only at certain time points ^41^ (*Table 3*). It has been suggested that muscle damage-attracted macrophages clean tissue via phagocytosis, which in turn convert macrophages into healing macrophages driving myoblast fusion and growth, fibrosis, vascularization, and return to homeostasis^42, 43^. TNF-α and IL-1β induce the proliferation and differentiation of myoblasts^44^. IL-1β decreases the level of myostatin, a negative regulator of muscle growth ^45^. Ablation of TNF-α or IL-6 displays poor muscle regeneration^37^. The CCR2-CCL2 pathway can promote macrophage recruitment, insulin-like growth factor-1 (IGF-I) production and stimulation of muscle regeneration^46^. TLR4 knock-out mice develop severe muscle injury^47^. Importantly, mild inflammation with low TNF-α and scarce macrophage infiltration can lead to poor muscle regeneration^48^. Our results are consistent with these multiple findings pointing to an important role of escalating inflammation in muscle recovery from injury and resolution of TMDM, while mild inflammation creates a negative environment for muscle regeneration. It was also suggested that M2 macrophages may stimulate myogenic precursor cell commitment into differentiated myocytes and the formation of mature myotubes^49^. However, our data indicated that IL-10 and IL-4 were not upregulated in MM by CFA, 0.2U and 10U Col at any time point.

In conclusion, based on our findings, we favor the hypothesis that escalating inflammation in MM is critical to resolve orofacial hypersensitivity triggered by tissue damage mimicked by Col intra-MM injection. Our results advance our understanding of the mechanisms controlling TMDM chronicity and provide an informatic basis for further studies on the regulation of gene expression plasticity in muscle affected by myalgia. The molecular basis of mechanisms controlling chronicity is only starting to emerge from preclinical and clinical studies but there is already evidence that it involves differences in cell types of the immune system.

## Ethical approval and informed consent

All experimental protocols were approved by the UTHSCSA IACUC committee. Protocol numbers are 20190114AR, 20220064AR and 20220069AR.

## Supporting information

Veh vs 1d post-CFA

Veh vs 1d post-Col 0.2U

Veh vs 1d post-Col 10U

Veh vs 5d post-CFA

Veh vs 5d post-Col 0.2U

Veh vs 6d post-Col 10U

Veh vs 14d post-Col 10U

## Acknowledgements

We would like to thank Mrs. Dawn Garcia and Mrs. Korri Weldon for assistance in the performance of the RNA-seq experiments; and Dr. Gregory Dussor (UTD, Dallas, TX) for advice on the behavioral tests. RNA-seq experiments were conducted in the Genome Sequencing Facility (GSF) in the Greehey Children’s Cancer Research Institute (GCCRI) of UTHSCSA. The GSF facility has been constructed in part with the support from UT Health San Antonio, NIH/NCI P30 CA054174 (Cancer Center at UT Health San Antonio), NIGMS/NIH S10 Shared Instrumentation Grant Program (SIG) (S10OD021805-01 to Z.L.), and Cancer Prevention Research Institute of Texas (CPRIT) Core Facility Award (RP160732). The Flow Cytometry Shared Resource at UT Health San Antonio is supported by a grant from the National Cancer Institute to the Mays Cancer Center (P30CA054174), a grant from the Cancer Prevention and Research Institute of Texas (CPRIT) (RP210126), a grant from the National Institutes of Health (S10OD030432), and support from the Office of the Vice President for Research at UT Health San Antonio. This research work was supported by HEAL Initiative NIDCR/NIH DE029187 (to A.T. and A.N.A.) and the National Institute Of Arthritis And Musculoskeletal And Skin Diseases of the National Institutes of Health (NIH/NIAMS) through the NIH HEAL Initiative (https://heal.nih.gov/) The Restoring Joint Health and Function to Reduce Pain (RE-JOIN) Consortium UC2 AR082195 (to A.N.A.).

## Author Contributions

K.A.L, A.H.H., J.M., and S.A.S.: *methodology, investigation, visualization*. K.A.L, S.A.S., Y.Z., Z.L., A.T., and A.N.A.: *analysis, conceptualization*. Z.L., A.T., and A.N.A.: *research design*. K.A.L., A.T., and A.N.A.: *resources, supervision, funding acquisition, manuscript preparation*. All authors reviewed the manuscript.

## Data and Material Availability

RNA-seq data has been deposited to GEO Accession, the number is GSE190183. Supplementary excel files show the raw gene readings/counts per gene for all our sequencing experiments and RPKM data for each sample. These supplementary files are “*Veh vs 1d post-CFA (MM; G-L columns are RPKM)*” for RNA-seq from male mouse MM at 1d post-CFA intramuscular; “*Veh vs 5d post-CFA (MM; G-L columns are RPKM)*” for RNA-seq from male mouse MM at 5d post-CFA intramuscular; “*Veh vs 1d post-Col 0.2U (MM; G-O columns are RPKM)*” for RNA-seq from male mouse MM at 1d post-Col (0.2U) intramuscular; “*Veh vs 5d post-Col 0.2U (MM; G-O columns are RPKM)*” for RNA-seq from male mouse MM at 5d post-Col (0.2U) intramuscular; “*Veh vs 1d post-Col 10U (MM; G-O columns are RPKM)*” for RNA-seq from male mouse MM at 1d post-Col (10U) intramuscular; “*Veh vs 6d post-Col 10U (MM; G-N columns are RPKM)*” for RNA-seq from male mouse MM at 6d post-Col (10U) intramuscular; “*Veh vs 14d post-Col 10U (MM; G-O columns are RPKM)*” for RNA-seq from male mouse MM at 14d post-Col (10U) intramuscular injection.

## Additional Information

All authors declare that they have no competing interests.

---

**Supplementary Figure 1:**
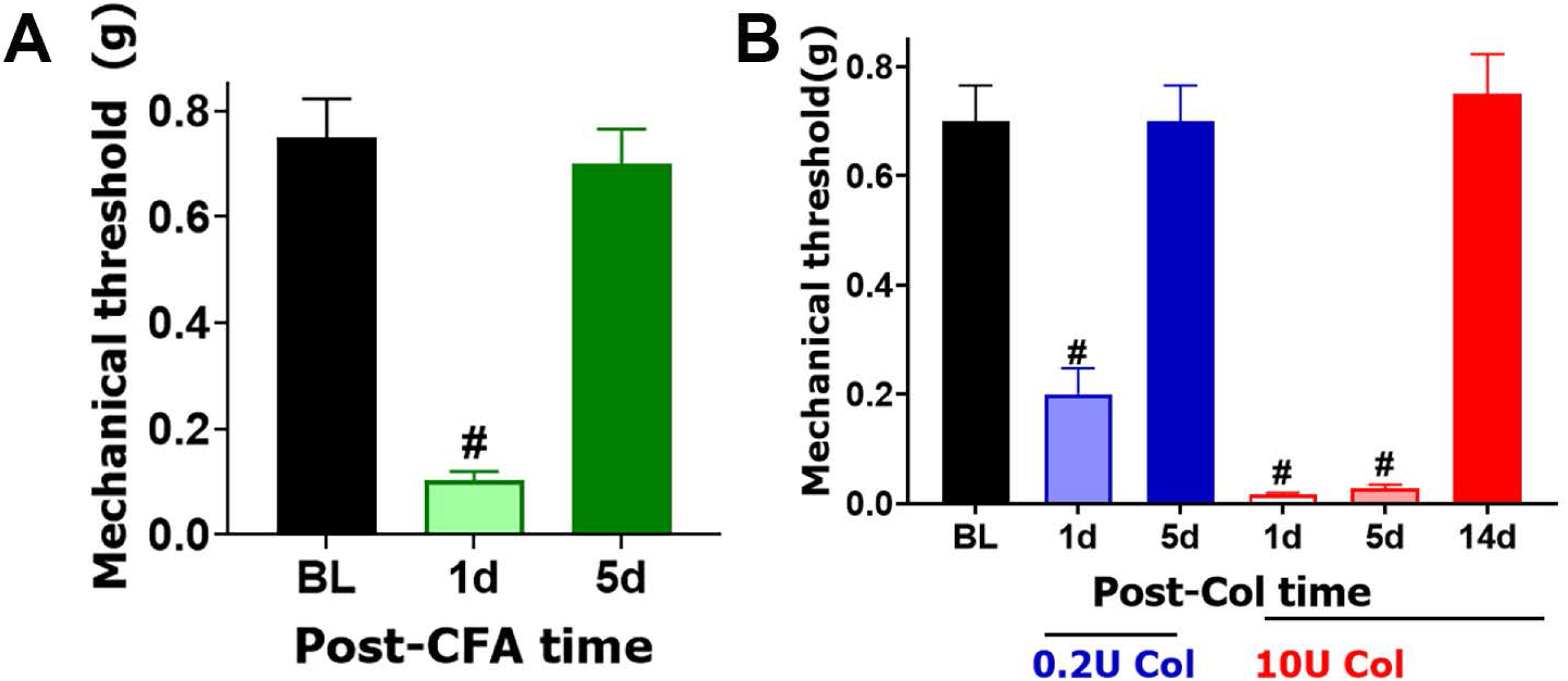
Col- and CFA-induced mechanical hypersensitivity in male mice. (**A**) Mechanical hypersensitivity at 1d and 5d post-CFA injected intro-MM in male mice. Statistical analysis is 1-way ANOVA Bonferroni’s post-hoc test (# p<0.0001; n=8). (**B**) Mechanical hypersensitivity at 1d and 5d post-0.2U Col and 1d, 5d and 14d post 10U Col injected intro-MM in male mice. Statistical analysis is 1-way ANOVA Bonferroni’s post-hoc test (# p<0.0001; n=8). X-axis indicates time post injection and injected reagent. Y-axis shows mechanical threshold assessed by von Frey filaments in an orofacial area above MM using up-down approach.

**Supplementary Figure 2:**
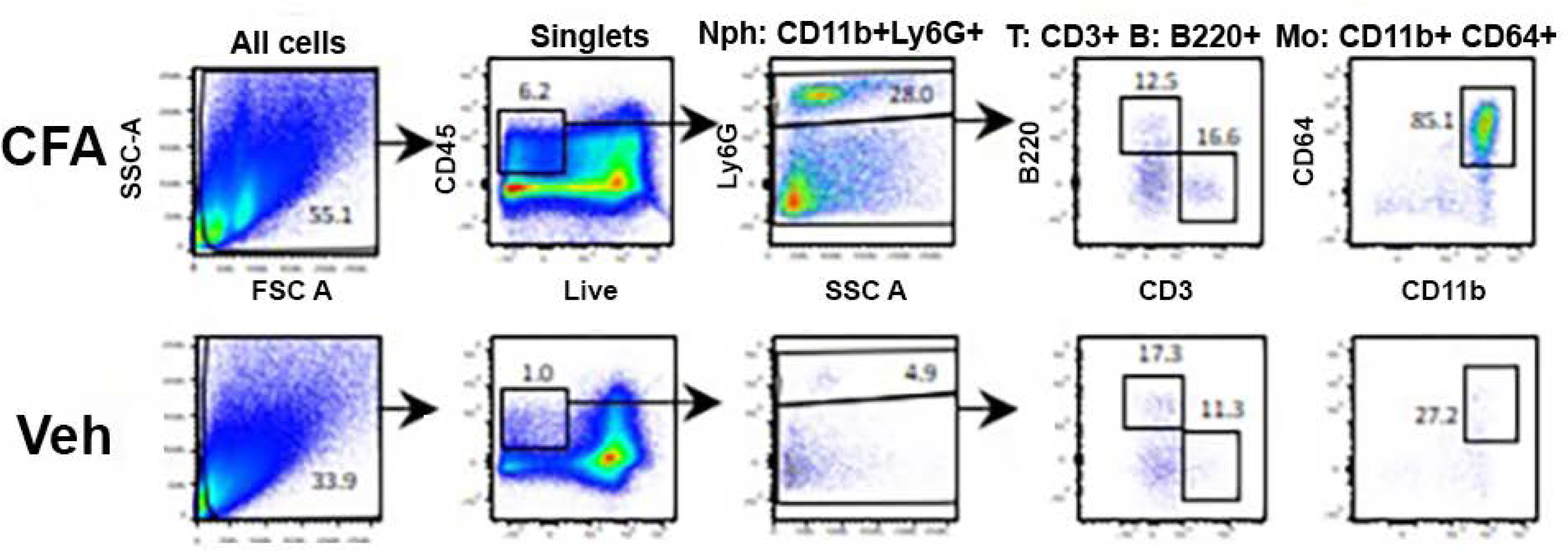
Raw data on gating strategy for flow cytometry. Gating strategy for single-cell suspension from the MM at 1d post-vehicle (Veh) or CFA single intramuscular treatment. The strategy shows live singlets and different immune cells were gated from single-cell suspension. Abbreviations are SSC-A - side angle scattered area; FSC-A - forward angle scattered area; CD45^+^ - immune cells; Mo - monocytes, Nph – neutrophils, CD3 – T-cells.

**Supplementary Figure 3:**
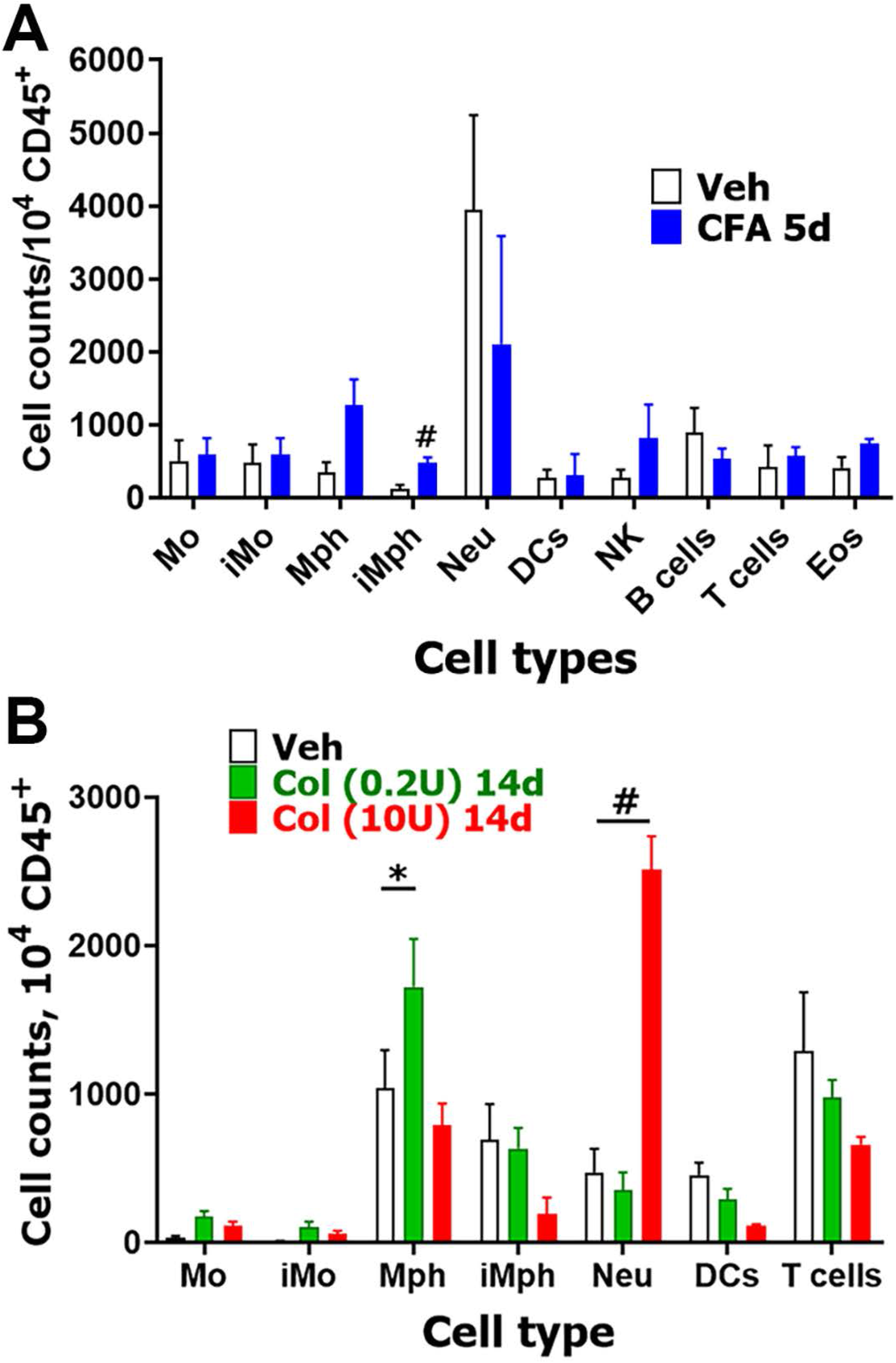
Immune cell profiles in CFA and Col treated MM at post-hypersensitivity resolution time points. **(A)** Immune cell counts per 10^4^ CD45^+^ cells in male MM at 5d post-vehicle (Veh) or CFA single intramuscular treatment. (**B**) Immune cell counts per CD45^+^ cells in MM at 14d post-vehicle (Veh), Col (0.2U) or Col (10U) single intramuscular treatment. DCs – dendritic cells; Mph – macrophages; iMph - inflammatory macrophages; Mo – monocytes; iMo - inflammatory monocytes; NK – natural killer cells; B – B-cells; T – T-cells and Neu – neutrophils. Statistic is 2-way ANOVA (* p<0.05; # p<0.0001; n=3-6).

